# 3D Reconstruction Enables High-Throughput Phenotyping and Quantitative Genetic Analysis of Phyllotaxy

**DOI:** 10.1101/2024.10.03.616344

**Authors:** Jensina M. Davis, Mathieu Gaillard, Michael C. Tross, Nikee Shrestha, Ian Ostermann, Ryleigh J. Grove, Bosheng Li, Bedrich Benes, James C. Schnable

## Abstract

Differences in canopy architecture play a role in determining both the light and water use efficiency. Canopy architecture is determined by several component traits, including leaf length, width, number, angle, and phyllotaxy. Phyllotaxy may be among the most difficult of the leaf canopy traits to measure accurately across large numbers of individual plants. As a result, in simulations of the leaf canopies of grain crops such as maize and sorghum, this trait is frequently approximated as alternating 180° angles between sequential leaves. We explore the feasibility of extracting direct measurements of the phyllotaxy of sequential leaves from 3D reconstructions of individual sorghum plants generated from 2D calibrated images and test the assumption of consistently alternating phyllotaxy across a diverse set of sorghum genotypes. Using a voxel-carving-based approach, we generate 3D reconstructions from multiple calibrated 2D images of 366 sorghum plants representing 236 sorghum genotypes from the sorghum association panel. The correlation between automated and manual measurements of phyllotaxy is only modestly lower than the correlation between manual measurements of phyllotaxy generated by two different individuals. Automated phyllotaxy measurements exhibited a repeatability of *R*^2^ = 0.41 across imaging timepoints separated by a period of two days. A resampling based genome wide association study (GWAS) identified several putative genetic associations with lower-canopy phyllotaxy in sorghum. This study demonstrates the potential of 3D reconstruction to enable both quantitative genetic investigation and breeding for phyllotaxy in sorghum and other grain crops with similar plant architectures.

## Introduction

Increases in crop productivity and water use efficiency are required due to increases in the world population and decreasing access to fresh water for agriculture [1]. In the past, increasing the tolerance of crops to high planting densities has improved crop productivity [2].

The increased tolerance of high planting densities in modern maize hybrids is explained at least in part by a shift in the distribution of light throughout the canopy [2], a distribution determined by plant canopy architecture. Photoinhibition in the upper canopy is decreased, and the photosynthetic capabilities of the leaves in the lower canopy are more effectively utilized when light is distributed more evenly throughout the canopy, increasing the overall radiation use efficiency of the crop [3]. Furthermore, shifting a larger proportion of photosynthesis into the lower canopy reduces the water loss via transpiration. Stomata lower in the canopy are less exposed to wind and thus have a stronger boundary layer; the water concentration gradient driving transpiration is additionally decreased as a consequence [4].

Interest in optimizing crop canopy architecture has motivated the study of genes and genomic loci determining variation in many of the individual components of canopy architecture, including the vertical leaf angle [5–8], internode length [9–12], and plant height [13–16]. Phyllotaxy is the arrangement of leaves around the stem in the horizontal plane (see Figure 1), and it also contributes to light distribution throughout plant canopies. Extreme phyllotaxic deviations and their inheritance have long fascinated geneticists [17]. However, relative to other canopy architecture traits, phyllotaxy has been subject to comparatively fewer quantitative genetic investigations. This absence may be explained, at least in part, by the difficulty of collecting large numbers of accurate measurements of phyllotaxy manually. As a result of the limited investigation of this trait, it has been unclear how much, if any, quantitative genetic variation in phyllotaxy exists in grain crops relative to the expectation of perfectly alternating – 180° degree angles between sequential leaves – phyllotaxy for these species.

**Figure 1.**
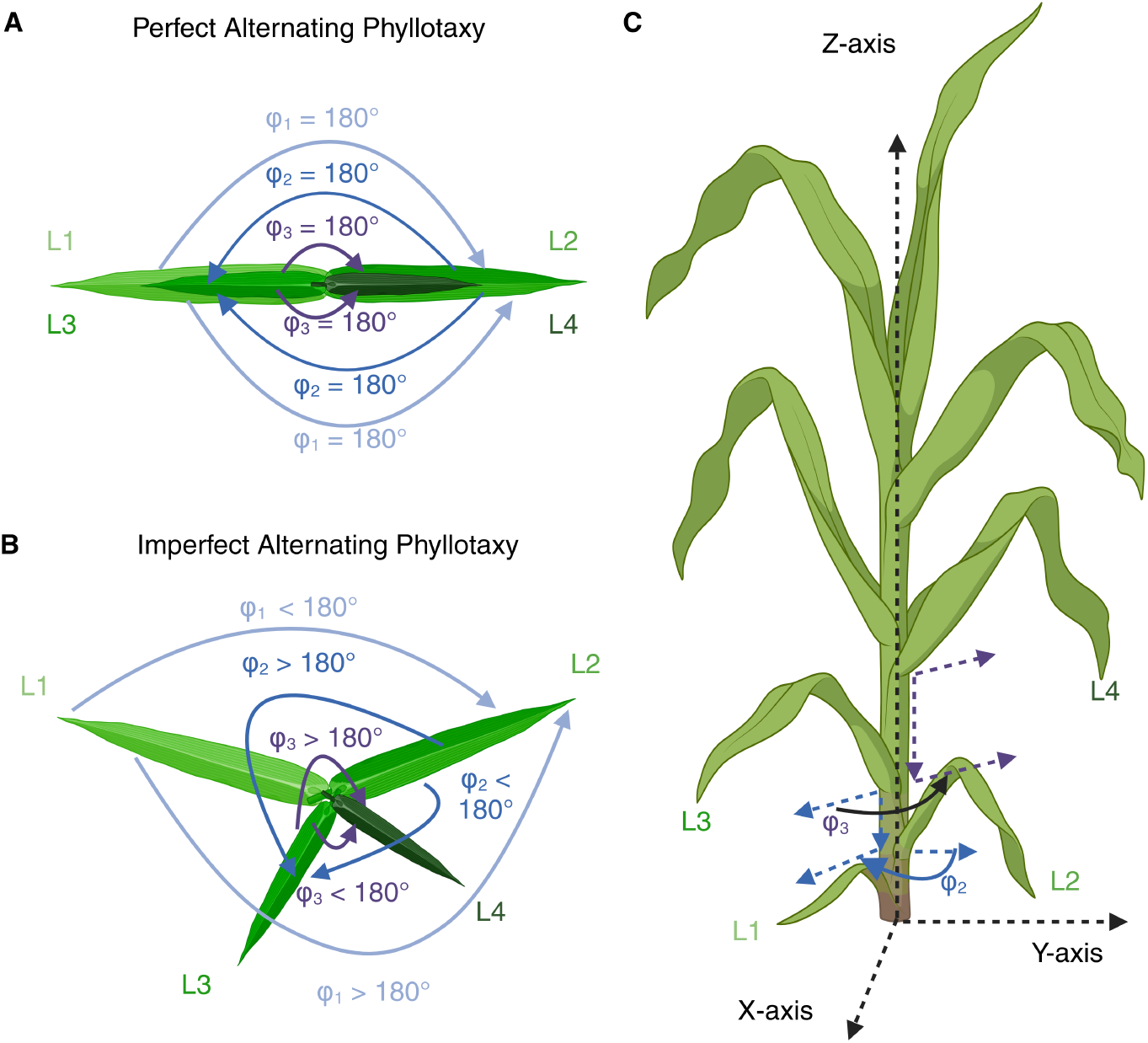
Phyllotaxy is the arrangement of the leaves around the stem in the *xy* plane. Created in Biorender.com [18]. **A)** The top view of a sorghum plant exhibiting the expected alternating phyllotaxy with 180° angles between each pair of sequential leaves. **B)** The top view of a sorghum plant exhibiting deviations from the expected phyllotaxic angles. Note that the angle captured will vary depending on the side of the plant measured. The two possible angles measured for a given pair of sequential leaves are conjugate to each other, *e*.*g*., their sum is equal to 360°. **C)** The side view of a sorghum plant and two examples of the phyllotaxic angle *φ* in the *xy* plane.

On a developmental level, phyllotaxy (Figure 1) is initially determined by the spacing of the newest leaf primordium, P0, in the shoot apical meristem (SAM), relative to the previous leaf primordium. The first molecular markers of the development of a new leaf primordium are an auxin maximum around the point of the new leaf primordium formed by PIN1 convergence and a subsequent down-regulation of *KNOTTED-LIKE HOMEOBOX (KNOX)* genes [19]. However, the final orientation of mature leaves appears to also be under a degree of environmental control. A range of environmental factors influence the orientation of leaves in maize, including wind, planting density, seed orientation, and water stress [20–25]. Some, but not all, maize genotypes have also exhibited the ability to reorient the axis of their leaves to avoid overlaps between neighboring plants [21, 25, 26]. Specific genes and genomic loci governing variation in this capacity have been mapped via GWAS [25]. However, this reorientation typically shifts the orientation of leaves on both sides of the plant reciprocally rather than modifying the alternating pattern of phyllotaxy that is typically exhibited by maize and other related plant species. Genetic variation and control of mean phyllotaxy has not been evaluated in depth via quantitative genetic methods, although several large effect single gene mutations that alter phyllotaxy have been characterized in maize [27, 28].

Perhaps the best-known of these phyllotaxy mutants is the recessive *abphyl1* mutant in maize, described by Jackson and Hake in 1999 [27]. This mutant typically exhibits an opposite phyllotaxy wherein one node produces two leaf blades with the midribs separated by approximately 180°, with occasional switches to the wild-type pattern of alternate phyllotaxy occurring partway through growth following an intermediate transition node wherein two leaf blades adjacent to one another are partially fused [27]. The authors also describe alternate phenotypes of the mutant wherein the shoot splits into two shoots with alternate phyllotaxy or a dwarfed plant with what appeared to resemble spiral phyllotaxy. Giulini et al. [29] cloned the gene underlying this mutant and found it a cytokinin-inducible response regulator controlling SAM size. The described opposite or spiral phenotypes of the *abphyl1* mutant in comparison to the wild-type alternate phyllotaxy may predispose us to conclude that phyllotaxy only varies qualitatively, but not quantitatively.

Previous methods of quantitatively measuring phyllotaxy can largely be divided into purely manual methods, top down imaging based methods, and approaches based on 3D reconstruction (Figure 1A). Protocols for manual measurements include the use of a circular protractor to measure the change in angle between sequential leaves [30], using a compass aligned with leaf midribs to measure the angle of each leaf with respect to magnetic north [22], a wooden panel marked with angles [21, 26, 31], or simple visual assessment of the angle of individual leaves relative to the axis of planting [24, 32]. These approaches tend to be relatively low throughput, with measurements collected from dozens to hundreds of plants representing less than ten genotypes per experiment. Top down imaging, whether from a UAV or from an elevated ground based camera, can increase the throughput of phyllotaxy measurements [23, 25, 32, 33]. Estimates of phyllotaxy can be obtained from these top-down images through a range of approaches including manual scoring [23, 25], fitting bounding boxes to individual leaves [33] or detecting the positions of midribs [32]. These methods are limited to measurements of the azimuth angle in the upper canopy, which is the deviation of the leaves from the row line (direction of planting). Other studies utilize electromagnetic 3D plant digitizers to reconstruct the plant *in silico* [30, 34]. While these methods offer precise measurements throughout the canopy, they are relatively low-throughput, as each plant required approximately 20 minutes of labor [32]. Daviet et al. [35] utilize skeletonized 3D reconstructions of maize to measure the azimuth angle, in a method similar to the one presented here, but do not evaluate the efficacy of the method for the measurement of phyllotaxy.

We use 3D reconstructions of sorghum plants from a diversity panel [36] to enable high-throughput phenotyping of phyllotaxy in the lower canopy and identify genetic markers associated with this trait. We identify heritable variation in sorghum phyllotaxy as well as three genetic markers associated with the median phyllotaxic angle in the lower 5 leaves. Application of this method to larger populations with additional replication will likely increase the number of marker trait associations for phyllotaxy in sorghum, providing a basis for both functional characterization of candidate genes and marker assisted selection.

## Materials and Methods

Figure 2 shows an overview of the workflow employed in this study. We describe each step in more detail below, but briefly: photos were taken from six different views (five side views and one top view) of 366 plants at three timepoints. The images were used as input to a 3D voxel carving algorithm described in [37]. The 3D voxels were skeletonized and segmented into the stem and leaves. Leaf angles are extracted by measuring the principal directions of the stem and leaves. We then normalized the angles, transforming them into the same coordinate system, and determined the angle difference in the *xy*-plane between successive leaves to estimate phyllotaxy. These values were then used to estimate the heritability of automated phyllotaxy measurements and conduct a GWAS analysis. Below, we describe each step in detail.

**Figure 2.**
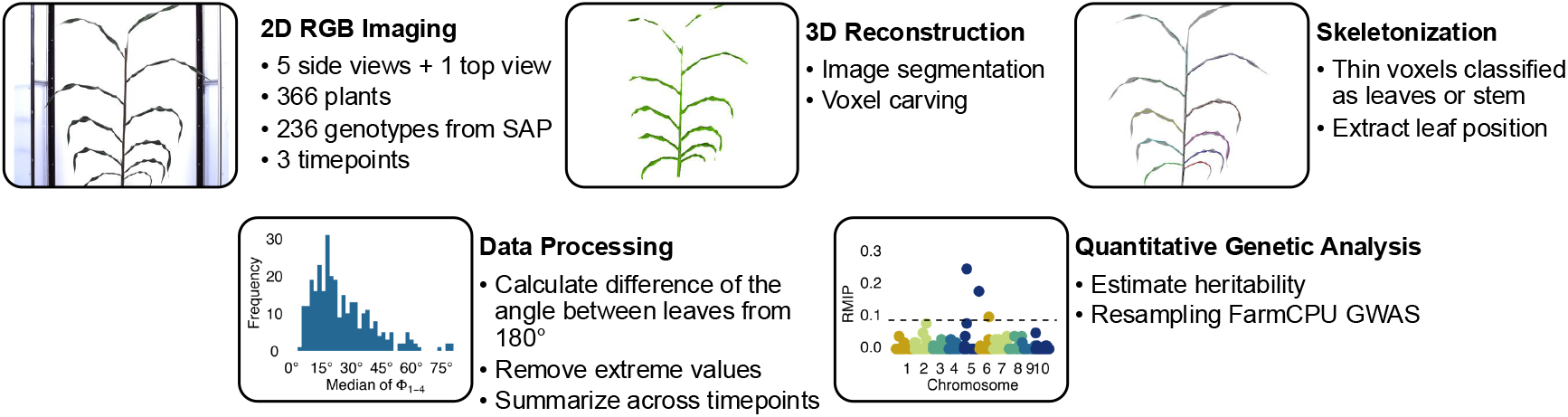
An overview of the workflow used in this paper and examples of output from each stage.

### Plant growth conditions, image acquisition, and manual measurements

A total of 366 sorghum plants, representing 236 genotypes from the sorghum association panel [36] with partial replication (40 replicated genotypes), were grown at the automated phenotyping facility of the University of Nebraska-Lincoln and imaged on April 11th, April 13th, and April 16th, 2018 (47, 49, and 52 days after planting, respectively). The growth and imaging protocols were followed as described in Tross et al. [5]. In 2023, an additional set of 10 sorghum plants were grown at the same facility and imaged on February 1, 2023, 76 days after planting. In 2024, a third set of 10 sorghum plants was grown at the same facility and manually measured and imaged on March 5, 2024, 47 days after planting. Between 2018 and 2023, the RGB camera used at the facility was upgraded from a Basler pia2400-17gc camera equipped with a c6z1218m3-5 Pentax TV zoom lens to a Prosilica GT6600 camera to improve resolution (from 2,454*×*2,056 pixels to 4,384*×*6,576 pixels) and image quality.

Manual phyllotaxy measurements were conducted by using the Compass application on either an iPhone 13 Pro Max or iPhone 14 [38] to measure the direction of each measured leaf. Differences between sequential leaves were calculated as was done for the automated phyllotaxy measurements. The left edge of the long side of the iPhone was aligned with the midrib of each leaf of interest and the phone was rotated along the *z*-axis (Figure 1C) until the short side of the screen was flush to the stem. This process was performed for each collared leaf on each plant of interest. Measurements of individual plants were repeated independently by two members of the research team to quantify the repeatability of manual phyllotaxy measurements.

### Phyllotaxy measurement from 3D skeletons

3D reconstructions of sorghum plants from 2D calibrated images were derived using methods described in Gaillard et al. [37, 39], and Tross et al. [5]. The images collected in 2018 were taken from five side views collected at equidistant angles (0°, 72°, 144°, 216°, 288°) around the plant. In 2023 and 2024, side view images were collected at 10 equidistant side views (0°, 36°, 72°, 90°, 108°, 144°, 216°, 252°, 288°, 324°). The images were calibrated to correct for potential misalignment between the pots and the turntable’s axis of rotation, as well as the camera’s optical center with the rotation axis. The calibrated images were then processed using a voxel carving algorithm producing a 512^3^ voxel resolution representation of the sorghum plant [37]. A skeletonization algorithm was applied to this voxel representation of the sorghum plant, iteratively removing voxels from the plant until only the skeleton structure remained. To eliminate any gaps in the skeleton caused by disconnected components, a joining process was implemented as part of the skeletonization process [39].

A Support Vector Machine (SVM) classifier [40] was employed to identify and discard portions of the skeleton that did not correspond to actual plant organs, for instance, spurious branches present from noise in the data. Post-processing techniques were used to classify voxels as either leaf or stem by computing paths from the ground to the leaves and labeling a voxel as part of the stem if it belonged to more than two paths. The separated leaves were then assigned numerical labels based on the attachment height to the stem within the skeleton structure. Using Principal Component Analysis (PCA), the stem voxels’ first and second principal axes were computed. Additionally, PCA was used to extract the principal axes for the first 20 voxels (about 6 cm) of each leaf, starting from the junction of the leaf with the stem. By using these two coordinate frames, the angles 0° ≤ *θ* < 180° and 0° ≤ *ϕ* ≤ 360° of each leaf were calculated in the stem coordinate frame, given by the PCA principal directions 3.

Accuracy and topology correctness were assessed as described in detail in Gaillard et al. [37, 39]. In brief, the accuracy scores were determined using the Dice coefficient based on the proportion of plant pixels in the 2D images represented by re-projected voxels in the 3D reconstruction. A topology was considered to be incorrect if the final plant skeleton did not exhibit a tree topology. Reconstructions with an accuracy score below 0.70 or an incorrect topology were removed from the dataset. Reconstructions containing only one *ϕ* value (i.e., only one leaf was identified in the skeleton) were also removed from downstream analyses as phyllotaxy relies on the angle between two leaves. This filtering criteria resulted in the exclusion of 115 reconstructions. Next, the differences between sequential leaves’ *ϕ* values were represented as *φ*_*i*_ = *ϕ*_*i*+1_ *− ϕ*_*i*_. Each *φ* value was then normalized to a range of *φ* ∈ [0°, 360°) by applying the modulo of 360.

### Method reliability measures and validation

#### Reliability of 3D-reconstruction and manual measurements

The lower five *φ* values generated for a plant from the 3D reconstructions were compared pair-wise between the three days of imaging (April 11th, April 13th, and April 16th, 2018) to estimate the reliability of 3D-reconstruction measurements of phyllotaxy. As it is rare for a healthy sorghum plant to have an angle less than a right angle and angles less than 90° could result from a leaf being missed during reconstruction or skeletonization, *φ* values less than 90° degrees or greater than 270° were removed.

There is no inherent structural difference between a phyllotaxy leaf angle of 160° or 200° with which of these two angles is reported by our method depending solely on the side of the plant measurement begins upon. To remove the arbitrary effect of side of the plant in comparisons between different measurements, we first determined if measurement began on the same side of the plant for both sets of measurements being compared. If the Pearson correlation coefficient on a per-plant basis was greater than or equal to zero, measurement began on the same side of the plant in both sets of measurements, and no transformation was applied to either set of measurements for that plant. In the cases where measurement began on different sides of the plant between the two sets of measurements for a given plant, we transformed the angles for this plant in one set of measurements (shown on the y-axes of plots in Figures 4 and S1) using:

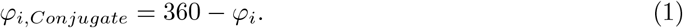

**Figure 3.**
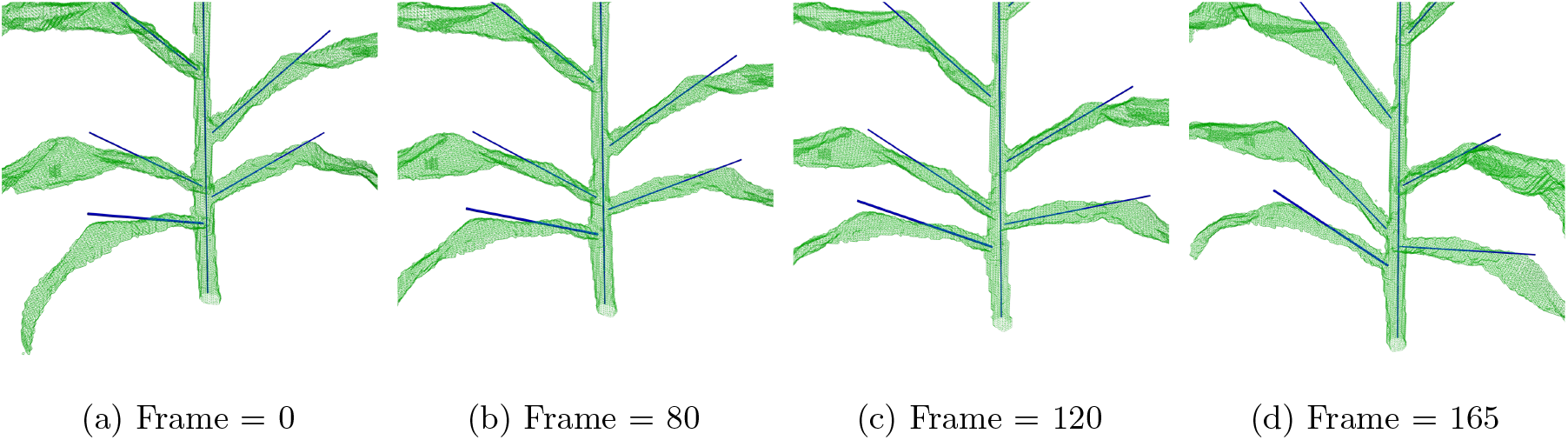
Rendering of a 3D reconstructed plant from the validation dataset, with the principal directions of the stem and leaves marked. The green wireframe shows the hull of the 3D reconstructed plant, and the solid blue lines show the principal directions of the stem and leaves of the plant. The supplemental material associated with this manuscript contains video animations showing the 3D reconstructions and leaves’ principal directions for each of the ten plants grown in 2023. We highly recommend the reader to watch the video animations to get a better sense of the angles in 3D.

**Figure 4.**
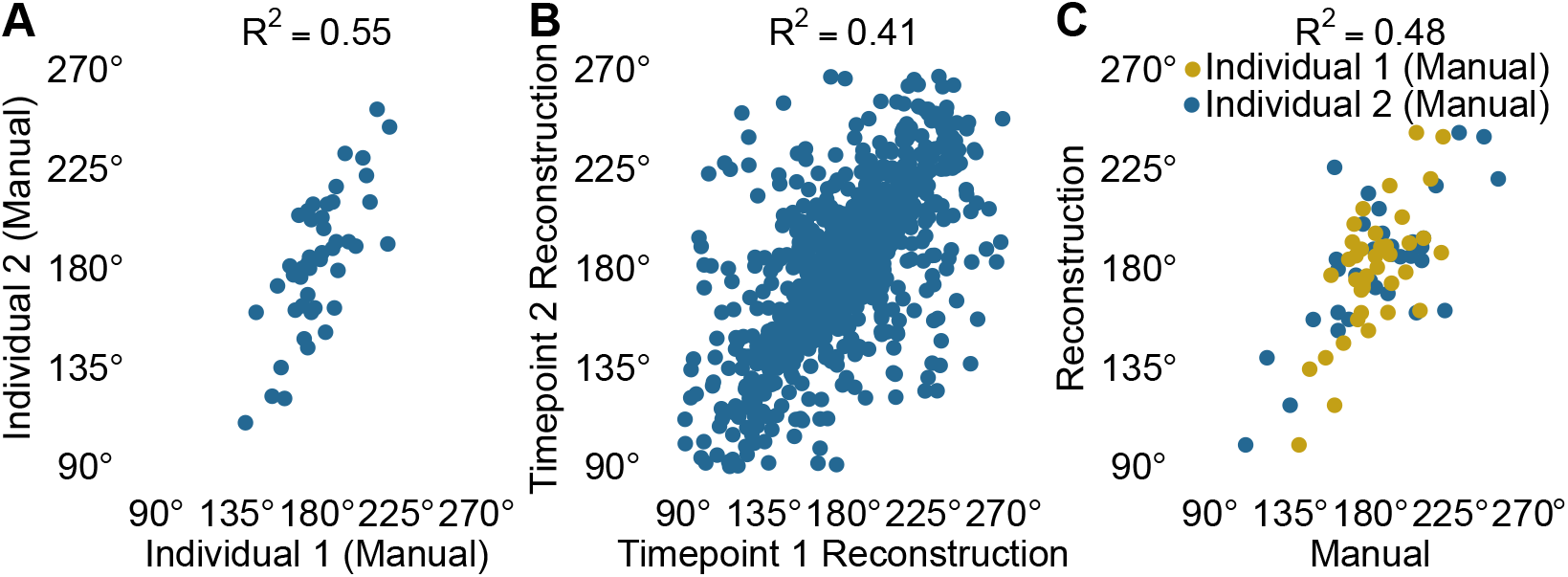
Both manual and reconstruction-based methods generate moderately repeatable measurements of phyllotaxic angles in the lower canopy. **A)** Correlation between manual measurements of lower five phyllotaxic angles (*φ*) for ten plants by two different people after removing *φ* values less than 90° or greater than 270°, with *R*^2^ = 0.55. **B)** Correlation of lower five *φ* values measured by 3D reconstructions of the sorghum plants when imaged on April 11, 2018 (Timepoint 1) and April 13, 2018 (Timepoint 2) after removing *φ* values less than 90° or greater than 270°, with *R*^2^ = 0.41. In cases when the measurements for a single plant were negatively correlated between the two days, the values from the Timepoint 2 reconstruction were transformed to their conjugate angles. **C)** Correlation between manual and 3D reconstruction measurements of lower five *φ* values after removing *φ* values less than 90° or greater than 270°, with *R*^2^ = 0.48. In cases where the 3D reconstruction and manual measurements were negatively correlated for a single plant, the 3D reconstruction measurements were transformed to their conjugate angles.

On a per-plant basis, this produces the same absolute value for the correlation between the two sets of measurements.

#### Comparison of 3D-reconstructions with manual measurements

The inital step of our voxel-carving and 3D reconstruction pipeline was segmentation of 2D input images into plant and non-plant pixels. For the 2018 images this step was performed using a convolutional neural network [37] trained specifically on manual annotations of the 2018 image dataset. For the 2023 and 2024 images taken with a different and higher resolution camera, we retrained a neural network with the same published architecture using 18 images collected from three plants imaged in 2023 and manually segmented using Paintbrush 2.6 [41] as well as an additional 67 images which were initially segmented using the older model trained on images from 2018 and then manually corrected and updated by human annotators. The segmented images output by the re-trained neural network were then used to reconstruct the sorghum plants. As the neural network was re-trained with images from the new camera, the reconstructions were visually checked. All 360°-normalized *φ*_*i*_ were obtained for the manual and 3D reconstruction measurements.

### Quantitative genetic analyses

For the purposes of quantitative genetic analysis, phyllotaxy values were transformed into the absolute difference between observed angle between two leaves and the expected angle for perfectly alternating phyllotaxy (180°): Φ, denoted as Φ_*i*_ = |*φ*_*i*_ *−* 180°|. Due to previous evidence [5] that heritability of measurements from the 3D reconstructions decreased at higher leaves due to movement of the upper canopy during rotation in the imaging process, we limited our analysis to the lower four phyllotaxic angles from five leaves. The 2D images of plants with a median Φ_*i*_ value greater than 1.5 times the interquartile range (136.1 *−* 167.0°) for one of the first 4 phyllotaxic angles were visually examined to determine if these images supported the extreme phyllotaxic values. In 55% of cases, the images could not definitively support the extreme phyllotaxic angles. Because it is rare for a healthy sorghum plant to have an angle less than a right angle and we could not verify the majority of extreme phyllotaxic angles visually examined, Φ_*i*_ *>* 90 were removed from downstream analyses. As lower-canopy phyllotaxy had not been extensively studied in previous literature, we evaluated 25 quantitative summaries of lower-canopy phyllotaxy to summarize across the three timepoints and/or multiple phyllotaxic angles, detailed in Supplementary Table 1. In aggregate, the criteria for reconstruction accuracy, skeleton topology, and Φ_*i*_ value resulted in the exclusion of 52 sorghum plants, leaving a total of 314 plants representing 223 unique genotypes for downstream quantitative genetic analyses. Data analysis and data visualization was conducted using R 4.2.2 [42] using the libraries lme4 [43], tidyverse [44], readxl [45], cowplot [46], MoMAColors [47], BiocIO [48], GenomicRanges [49], Gviz [50], ggrepel [51], scales [52], viridis [53], stringi [54], and car [55].

#### Heritability

The proportion of variance in each quantitative summary of phyllotaxy attributable to differences between genotypess, broad-sense heritability, was estimated using data from 40 genotypes represented by multiple plants in the filtered dataset. The linear model

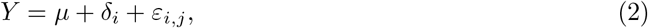

was fit using the R package lme4 [43], where *Y* is the response variable, *µ* is the overall mean, *d*_*i*_ is the random effect of the *i*th genotype, and *ε*_*i,j*_ is the residual error for the *j*th plant of the *i*th genotype. Variance components were then extracted, and broad sense heritability was calculated as

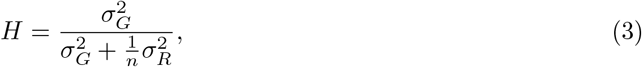

where 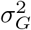 is the genotypic variance and 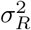 is the residual variance and *n* = 2, the minimum number of replications per genotype. When there were more than 2 replications per genotype, all replicates were used in the estimation except in the case of the reference genotype PI656058, which was replicated 8 times, 2 more than any other genotype. 2 random replications of PI656058 were used in the estimation.

#### GWAS

Genome-wide association studies reported here were conducted for the 13 quantitative summaries of phyllotaxy with a broad-sense heritability ≥ 0.20 from 218 sorghum varieties which were phenotyped as part of this study, passed the quality control steps described above, and were present in a set of 4,693,810 genetic markers scored for the same population via whole genome resequencing. Five of the 223 genotypes that passed the quality control steps described above were not present in the larger set of genetic markers called for the sorghum association panel [56] aligned to the Sorghum bicolor v3.1.1 reference genome [57] which was filtered to generate the genetic marker set used in this study. These five genotypes were excluded from the GWAS. The marker set from Boatwright et al. [56] was filtered to exclude those markers which were not biallelic, indels, or missing in ≥ 30%, heterozygous in *≥* 10%, or with a minor allele frequency <5% of the 218 genotypes present in both the phenotype and genotype data using VCFtools v0.1.16 [58] and BCFtools 1.17 [59]. The number of effective markers was estimated to be 1,088,251.19 using GEC v0.2 [60]. The dataset was analyzed one hundred times using the FarmCPU algorithm as implemented in rMVP v1.0.6 [61], with a threshold of 0.21 for iteration in the FarmCPU algorithm, corresponding to approximately 0.05 divided by the ratio of estimated effective markers to total markers. In each interation, 10% of phenotypic records were randomly masked and genetic markers with a p-value of less than 4.59 *×* 10^−8^, corresponding to a Bonferroni corrected p-value of 0.05 applied to the estimated effective independent genetic markers, were considered to be significantly associated with the phenotype. For each marker which exceeded this threshold in at least one of the one hundred iterations, a resampling model inclusion probability (RMIP) was calculated based on the number of interations in which the marker exceeded the significance threshold divided by the total number of interations. Markers that exceeded an RMIP of 0.1 were considered of greatest interest for downstream analysis. Linkage disequilibrium estimates within a chromosome were estimated using plink v1.90 [62].

## Results

### Reliability of automated 3D phyllotaxy measurements

Measuring each leaf angle for a single plant took one person between ten and twenty minutes to complete. After removing extreme values and transforming conjugate angles as described in methods, the correlation of ground truth measurements of phyllotaxy between different individuals measuring the same pairs of leaves on the same plants was *R*^2^ = 0.55 (*n* = 46 angles), based on data from five pairs of leaves per plant measured on ten plants by two individuals (Figure 4A). This was modestly higher than the correlation between measurements of the first five pairs of leaves generated from 3D reconstructed plants imaged at two time points separated by two days (*R*^2^ = 0.41, *n* = 961 angles, Figure 4B) after applying the same filtering criteria and transformations to the data. When the same comparison was made between automated measurements of phyllotaxy generated using images collected either three days apart or five days apart, the correlation between measurements declined to *R*^2^ = 0.33 and *R*^2^ = 0.24, respectively (Figure 4B, Supplementary Figure 1). We also found a moderate correlation (*R*^2^ = 0.48, *n* = 75 angles, Figure 4C) between 3D reconstruction measurements and manual measurements when comparing across the combined set of manual measurements taken by Individual 1 and Individual 2 and the corresponding conjugate angles from the reconstructions.

### Variation in sorghum phyllotaxy

An initial assessment of phyllotaxy in 336 plants (236 genotypes) of the sorghum association panel using the 3D reconstruction method described above identified plants with deviations from the expectation of perfectly alternating phyllotaxy between the second and third extant leaves in sorghum plants ranging from Φ_2_ = 1.05° (nearly perfectly alternating phyllotaxy between leaves 2 and 3, Figure 5A, D) to Φ_2_ = 170.4° (leaves 2 and 3 emerging one on top of the other, Figure 5C, F). Absolute variations from the expected angle of 180° in the lower 4 phyllotaxic angles across all plants and all timepoints ranged from Φ_*i*_ = 0.01° to Φ_*i*_ = 179.97 (Figure 5G). In some cases, extreme phyllotaxy values could be validated by manual examination of the source images (Figure 5B, C, E, F). However, in 55% of cases visually examined, manual examination of source images for sorghum plants that deviated from the expected phyllotaxy by >90° could not definitively support these extreme values. In some cases, specific issues were identified to which the incorrect measurements could be attributed including the presence of one or more tillers, fallen plants, or leaves senescing in an unexpected order (Supplemental Figure 2), as well as errors in the ordering of leaves. Given the difficulty even trained subject matter experts experienced in accurately assessing phyllotaxy from 2D images and the high rate of errors among manually checked phyllotaxy angles in the *>*90° bin, the decision was made to exclude these values from downstream analysis.

**Figure 5.**
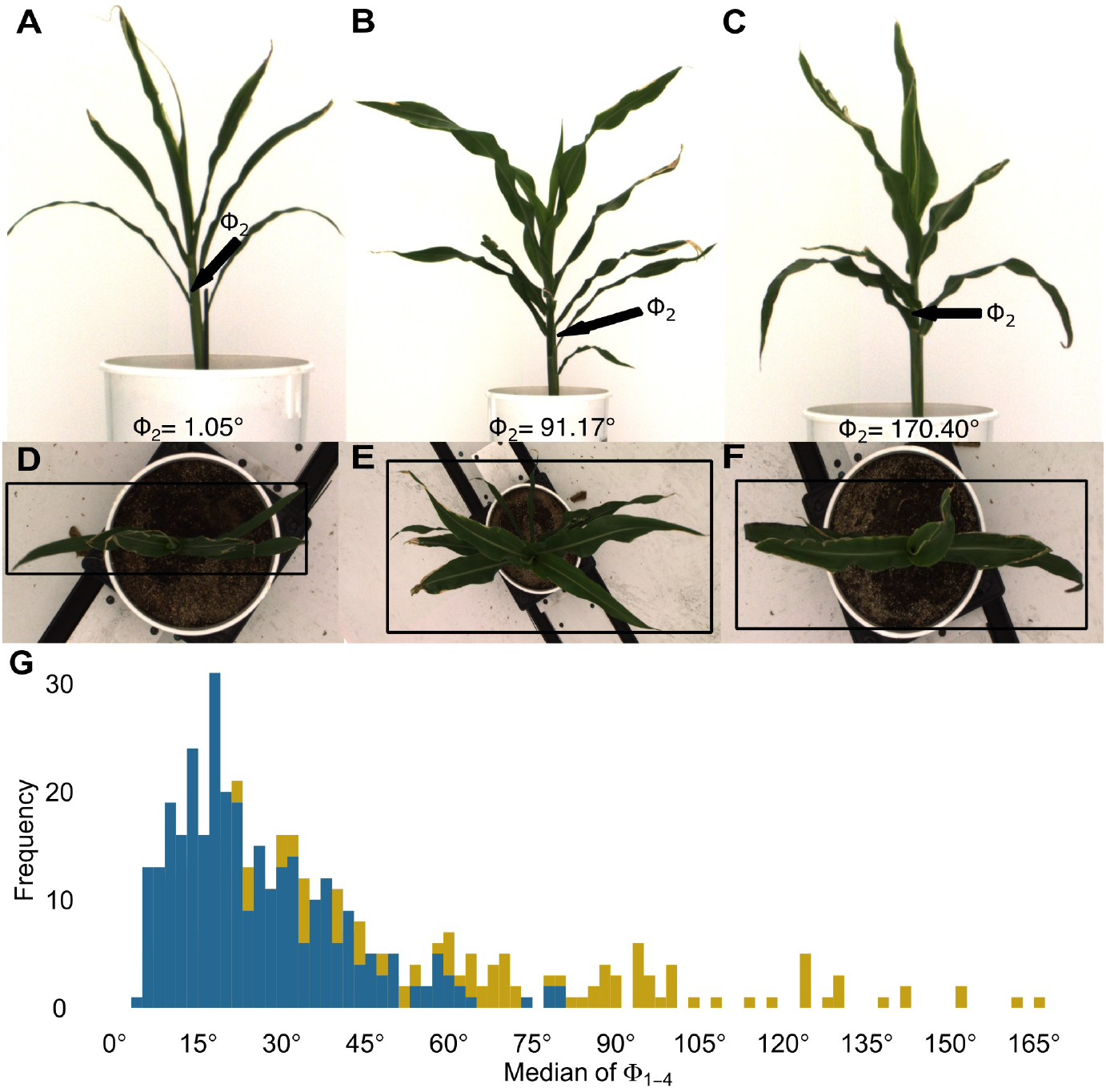
Variation of phyllotaxic angles among sorghum plants of different genotypes. **A-F)** Side (A-C) and top (D-F) views of three sorghum plants with minimal (A, D, PI533866), moderate (B, E, PI533852), and extreme (C, F, PI533915) levels of deviation from the expected value of 180° for Φ_2_.**G)** The distributions of median Φ_1*−*4_ values before (gold, 366 plants) and after removing extreme values (blue, 308 plants).

### Quantitative genetic analysis

As we had measurements for multiple phyllotaxic angles in the lower canopy per plant across the three timepoints, we evaluated 25 quantitative metrics of lower canopy phyllotaxy to summarise across multiple timepoints and/or angles after removing extreme values, which are described in Supplemental Table 1. Thirteen of 25 quantitative metrics were estimated to have broad-sense heritabilities greater than or equal to 0.2 (Supplemental Table 1). We detected stable (RMIP *≥* 0.1) marker-trait associations for 7 of these 13 quantitative summaries of lower canopy phyllotaxy via GWAS (Figure 6, Supplementary Table 1, Supplementary Figure 3). Six of the seven markers which exceeded an RMIP of 0.1 for at least one trait were also identified across four or more total traits when the RMIP threshold was reduced to 0.02. These six genetic markers and their associated traits are detailed in Supplementary Table 2. The repeated signals associated with the same markers indicate that multiple phyllotaxy summary metrics capture similar information content about the properties of the lower canopy in sorghum.

**Figure 6.**
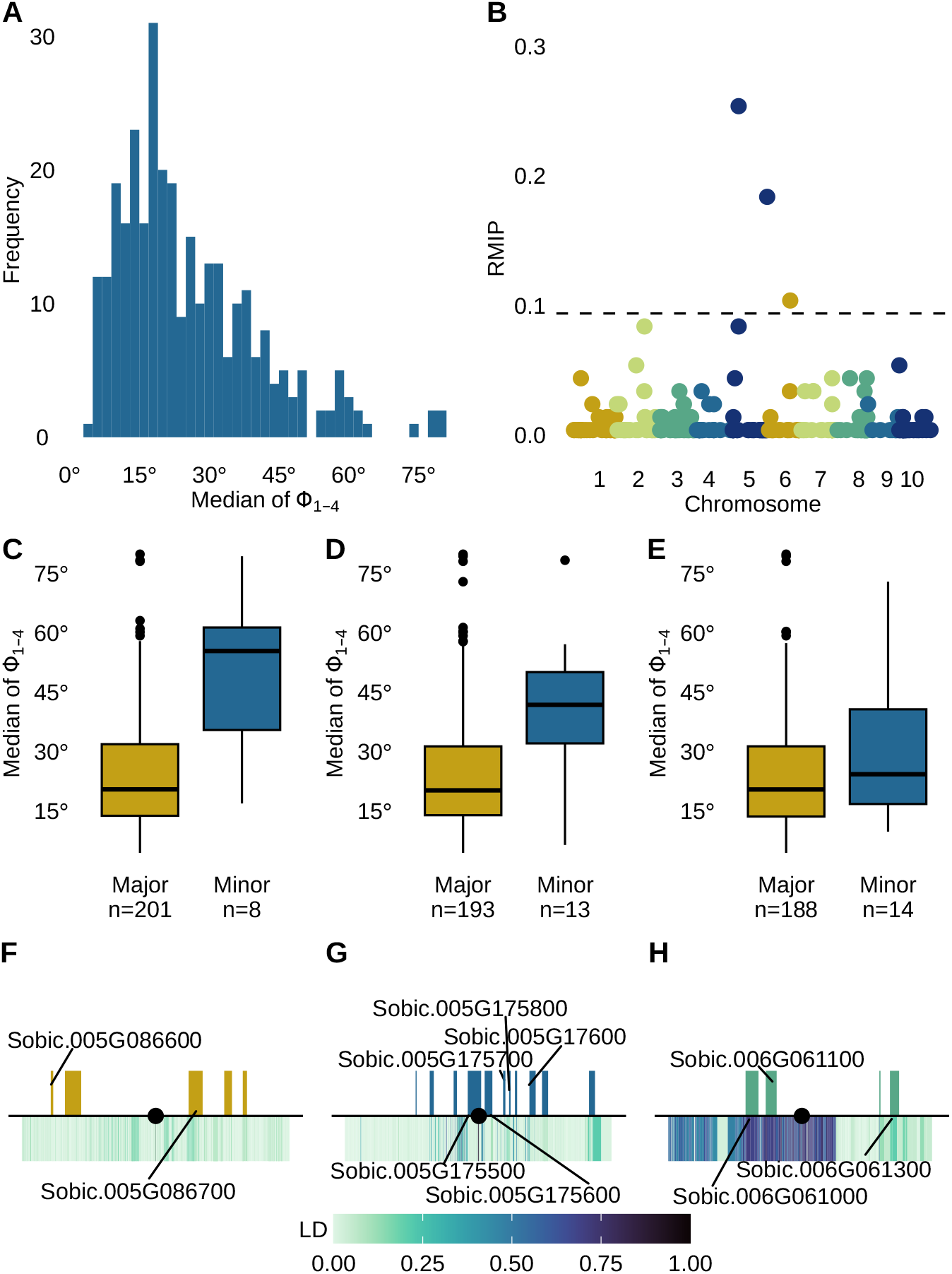
GWAS identifies genomic regions associated with variation in the median of the lower four phyllotaxic angles. **A)** Distribution of the median of the Φ_1*−*4_ values for each plant. The broad sense heritability of this trait was estimated to be 0.25. **B)** Results of a resampling FarmCPU GWAS conducted for median of the Φ_1*−*4_ value. Dashed line indicates an RMIP value of 0.10, the cutoff employed in this study. **C)** Median Φ_1*−*4_ values for each plant by allele (major or minor) at Chr05:12,109,370 (RMIP= 0.26). The *n* below each box indicates the number of genotypes homozygous for the allele. Genotypes with heterozygous calls at the marker were excluded. **D)** Median Φ_1*−*4_ values for each plant by allele (major or minor) at Chr05:65,733,791 (RMIP= 0.19). **E)** Median Φ_1*−*4_ values for each plant by allele (major or minor) at Chr06:41,390,777 (RMIP= 0.11). **F)** Genomic interval surrounding the trait associated marker Chr05:12,109,370 (black dot). The total region shown is 200 kilobases, 100 kilobases on either side of the trait associated marker. Colored boxes above the black line indicate the position of annotated genes. Color bar below the black line indicates linkage disequilibrium between the trait associated marker and other genetic markers within the 200 kilobase interval. **G)** Genomic interval surrounding the trait associated marker Chr05:65,733,791. **H)** Genomic interval surrounding the trait associated marker Chr06:41,390,777.

The marker-trait association with the highest stability was identified for the median Φ_1*−*4_ value with the genetic marker Chr05:12,109,370 (RMIP= 0.26, Figure 6). This trait captures the median phyllotaxic angle measured in the lower four phyllotaxic angles across all three timepoints for a plant, and had a relatively normal phenotypic distribution (Figure 6A). Two additional markers, Chr05:65,733,791 and Chr06:41,390,777, were also identified to have stable associations with this trait (RMIP= 0.19, 0.11, respectively; Figure 6B). Genotypes carrying the minor allele at each of these sites have significantly higher (*p <* 0.05) median deviations from the expected phyllotaxic angle of 180° than genotypes carrying the major allele at the same site (Figure 6C–E). Figure 6F–G shows the annotated gene models within 100 kb of the identified genetic markers in order of descending RMIP, and all annotated gene models within this region and their functional annotations are described in Supplementary Table 3. The nearest gene model to the first genetic marker on chromosome 5, Sobic.005G086700, which encodes a zinc finger transcription factor, is located 24.5 kb from the trait associated marker. Linkage disequilibrium (LD) decays quickly in this region, with the maximum LD between the identified genetic marker and genetic markers within the nearest gene model being *R*^2^ = 0.18. Within 100kb of this trait associated marker, there is also a gene encoding an O-methyltransferase (Sobic.005G086600) and three gene models with no functional annotation. The second trait associated marker on chromosome 5 is located in the sixth intron of Sobic.005G175500, which is annotated as encoding a protein tyrosine kinase related to salt stress and antifungal responses. The trait associated marker on chromosome 6 is in LD (*R*^2^ *>* 0.6) with genetic markers in two gene models, Sobic.006G061000 and Sobic.006G061100 (Figure 6G). The closer of these two, Sobic.006G061100 located 18.8 kb from the GWAS hit, encodes an AMP-activated protein kinase. The second encodes a protein belonging to the pentatricopeptide repeat (PPR) family.

## Discussion

We present a high-throughput method of measuring phyllotaxy in the lower canopy that achieves near-human repeatability. The imaging process for each plant requires little human intervention and can be completed in approximately 2 minutes, and the reconstruction and skeletonization steps each run in less than one minute, making it far more high-throughput than the manual method we employed which requires 10 – 20 minutes per plant to produce data with comparable repeatability. The more rapid methodology enables application to a large panel of plants. Daviet et al. [35] used a similar technique as presented here of reconstruction and skeletonization and show its ability to measure the azimuth positions of maize leaves. However, they do not extend their study to characterize the position of subsequent leaves relative to each other, a key factor in the process of leaf initiation, nor do they perform quantitative genetic studies to illuminate potential genetic mechanisms of this under-studied trait. We evaluate the reliability of the method we present here and find it to be moderately stable (*R*^2^ =0.41 for the same plants imaged on different days) after removing extreme values. This is modestly less repeatable than manual measurements (*R*^2^=0.55 for the same plants measured by different people). It is possible that a portion of the residual value in correlations of phyllotaxy measurements collected at different time points is associated with subtle changes in plant growth or the senescence of leaves, as the repeatability of automated phyllotaxy measurements decreases when comparing images collected with larger time intervals (see Supplementary Figure 1). We detect genetically repeatable variation in the lower 4 phyllotaxic angles of the canopy that is to some degree, robust to the specific summary metric used to summarise across timepoints and/or multiple angles. Thirteen of the 25 summary metrics were estimated to have broad-sense heritabilities greater than 0.20 (Supplementary Table 1). Several of these measures of lower-canopy phyllotaxy utilize information from multiple angles, while others include information only from a single angle across timepoints.

While the method we present here represents a large step forward in the ability to illuminate the genetic basis for phyllotaxy in sorghum, it is not without its limitations. First, this method is constrained to a greenhouse setting, limiting the ability to study the effects of environmental and management factors on crop phyllotaxy and the resulting productivity under field conditions with a true crop canopy. Secondly, the method has the highest accuracy in the lower canopy due to occlusion of the point of attachment by other leaves in the middle and upper canopy or significant movement of the upper plant during the imaging process, which makes accurate reconstruction difficult. While phyllotaxy in the upper canopy can be estimated using methods similar to He et al. [33], the middle canopy remains a challenge to measure, and evidence exists that different levels of the canopy have different genetic determinants of architecture [5, 6]. Third, the method discussed here may generate extreme values when tillers are present (Supplementary Figure 2). Fourth, the method is highly dependent on camera calibrations and high-quality image segmentations to train and improve the convolutional neural network to segment the RGB images that serve as input to the pipeline for this method. Fourth, this initial study was done using a relatively small population (218 genotypes) for association studies, limiting the statistical power available to detect genes controlling the trait of interest while sufficiently controlling for false positives.

Despite the limited statistical power provided by the small population employed, we identified marker-trait associations a total of 14 significant GWAS signals representing 7 unique markers linked to variation in one or more phyllotaxy summary metrics at RMIP*≥* 0.1(Figure 6, Supplementary Figure 3, Supplementary Table 1). Further examination of the marker-trait associations detected for the median Φ_1*−*4_ value, which had the most strongly supported marker trait association (RMIP= 0.26) showed that the genotypes with the minor allele at any of these sites have significantly higher median Φ_1*−*4_ values than genotypes with the major allele, indicating greater deviations from the expected alternating phyllotaxy of sorghum (Figure 6C–E). As rare alleles tend to be deleterious [63], this may indicate that deviations from alternating phyllotaxy are detrimental to overall fitness. The genomic regions surrounding the genetic markers associated with the median Φ_1*−*4_ value include several transcription factors, osmotic stress response genes, protein kinases, and calcium binding proteins, as well as several gene models with no functional annotations (Supplementary Figure 3). In the future, the method we present here could be employed to score phyllotaxy in larger association panels, such as the Sorghum Biodiversity Panel (SbDiv) [64] and/or to score more replicated plants per genotype, both of which should improve our statistical power to identify specific genomic intervals associated with variation in phyllotaxy. As maize and sorghum share highly similar plant architectures prior to the reproductive stage, we anticipate our method should also be applicable to this crop without the need for extensive modification or fine tuning. We demonstrate the feasibility of high-throughput measurements of lower-canopy phyllotaxy, enabling quantitative genetic analysis to improve our understanding of its genetic control.

## Supporting information

Figure 1 Publication License

## Data Availability

The code for reconstruction and skeletonization is available at GitHub: https://github.com/cropsinsilico/SorghumVoxelCarving

The raw images analyzed in this study are available at Zenodo: Mathieu Gaillard, Chenyong Miao, James C. Schnable, & Bedrich Benes. (2021). Voxel Carving Based 3D Reconstruction of Sorghum [Data set]. Zenodo. https://doi.org/10.5281/zenodo.4426620.

The phenotypic data, GWAS result files and code for main figures and analysis are available at Github: https://github.com/jdavis-132/phyllotaxy.git

## Author Contributions

JMD and NS collected measurements and ground truth data. MG IO and BL designed methods for and performed image analysis, plant reconstruction and trait value extraction. JMD NS and RJG annotated image data and employed domain expertise to reconcile extracted trait values and true plant architectures. JMD, BB, and JCS designed experiments and analytical approaches. JMD conducted statistical and quantitative analysis and visualized results with input from MCT and JCS. JMD drafted the manuscript with input from JCS and BB. All authors reviewed and approved the final version of the manuscript.

## Supplementary Materials

Supplementary File 1. Videos of reconstructed plants grown in 2023. https://github.com/jdavis-132/phyllotaxy/blob/f02e6b9ed52723510a78490d34153699266837c5/SupplementalVideos-Phyllotaxy.zip

**Supplementary Table 1:** Multiple quantitative summaries of phyllotaxy captured heritable variation.

| Summary of phyllotaxy | Description | Broad-sense heritability | Number genetic markers with RMIP $\geq 0.1$ |
| --- | --- | --- | --- |
| Median of $\Phi_{1-4}$ | Median of lower 4 phyllotaxic angles | 0.25 | 3 |
| Mean of $\Phi_{1-4}$ | Mean of lower 4 phyllotaxic angles | 0.30 | 5 |
| Mean of Median $\Phi_{1-4}$ | Mean of the medians of each of the lower 4 phyllotaxic angles | 0.27 | 2 |
| Median of $\Phi_{1-3}$ | Median of lower 4 phyllotaxic angles | 0.27 | 1 |
| Median of $\Phi_2$ | Median of the second phyllotaxic angle | 0.34 | 1 |
| Mean of $\Phi_2$ | Mean of the second phyllotaxic angle | 0.34 | 1 |
| Median of $\Phi_3$ | Median of the third phyllotaxic angle | 0.32 | 1 |
| Mean of $\Phi_{1-3}$ | Mean of the lower 3 phyllotaxic angles | 0.30 | 0 |
| Mean of $\Phi_3$ | Mean of the third phyllotaxic angle | 0.24 | 0 |
| Mean of Median $\Phi_{1-3}$ | Mean of the medians of each of the lower 3 phyllotaxic angles | 0.28 | 0 |
| Mean of Mean $\Phi_{1-3}$ | Mean of the means of each of the lower 3 phyllotaxic angles | 0.28 | 0 |
| Mean of Mean $\Phi_{1-4}$ | Mean of the means of each of the lower 4 phyllotaxic angles | 0.33 | 0 |
| Mean GLM BLUE Fitted $\Phi_{1-4}$ | Mean of fitted values for lower 4 phyllotaxic angles from GLM BLUE model | 0.28 | 0 |
| Mean of Means $\Phi_{1-2}$ | Mean of the means of each of the lower 2 phyllotaxic angles | 0.16 | |
| Mean of $\Phi_{1-2}$ | Mean of first 2 phyllotaxic angles | 0.15 | |
| Mean of Medians $\Phi_{1-2}$ | Mean of the medians of each of the lower 2 phyllotaxic angles | 0.14 | |
| Median of Means $\Phi_{1-4}$ | Median of the means of each of the lower 4 phyllotaxic angles | 0.05 | |
| Median of $\Phi_1$ | Median of first phyllotaxic angle | 0.03 | |

| Continuation of Table 1 |  |  |  |
| --- | --- | --- | --- |
| Summary of<br>phyllotaxy | Description | Broad-sense<br>heritabil-<br>ity | Number<br>genetic<br>markers<br>with<br>RMIP<br>$\geq 0.10$ |
| Median of Means<br>$\Phi_{1-3}$ | Median of the means of each of the lower 3<br>phyllotaxic angles | 0.02 | |
| Median of $\Phi_4$ | Median of fourth phyllotaxic angle | 0.00 | |
| Mean of $\Phi_1$ | Mean of first phyllotaxic angle | 0.00 | |
| Mean of $\Phi_4$ | Mean of fourth phyllotaxic angle | 0.00 | |
| Median of Medians<br>$\Phi_{1-3}$ | Median of the medians of each of the lower 3<br>phyllotaxic angles | 0.00 | |
| Median of Medians<br>$\Phi_{1-4}$ | Median of the medians of each of the lower 4<br>phyllotaxic angles | 0.00 | |
| Median of Means<br>$\Phi_{1-2}$ | Median of the means of each of the lower 2<br>phyllotaxic angles | 0.00 | |

**Supplementary Table 2:** Six genetic markers identified to have a stable association with one trait were sometimes identified to have associations with other traits.

| Chromosome | Position | Traits with RMIP<br>$\geq 0.10$ | Traits with RMIP<br>$\geq 0.02$ | Maximum<br>RMIP |
| --- | --- | --- | --- | --- |
| Chr05 | 12109370 | Median of $\Phi_{1-4}$,<br>Mean of $\Phi_{1-4}$,<br>Mean of Medians<br>$\Phi_{1-4}$ | Mean of Means<br>$\Phi_{1-4}$, Mean of<br>GLM BLUE Fitted<br>$\Phi_{1-4}$ | 0.26 |
| Chr05 | 65733791 | Median of $\Phi_{1-4}$,<br>Mean of $\Phi_{1-4}$,<br>Mean of Medians<br>$\Phi_{1-4}$ | Mean of Means<br>$\Phi_{1-4}$, Mean of<br>GLM BLUE Fitted<br>$\Phi_{1-4}$ | 0.19 |
| Chr03 | 56520108 | Median of $\Phi_2$,<br>Mean of $\Phi_2$ | | 0.12 |
| Chr06 | 41390777 | Median of $\Phi_{1-4}$,<br>Median of $\Phi_{1-3}$,<br>Median of $\Phi_3$ | Mean of $\Phi_{1-4}$,<br>Mean of $\Phi_{1-3}$,<br>Mean of $\Phi_3$, Mean<br>of Medians $\Phi_{1-3}$,<br>Mean of GLM<br>BLUE Fitted $\Phi_{1-4}$ | 0.11 |
| Chr05 | 5060741 | Mean of $\Phi_{1-4}$ | Median of $\Phi_{1-4}$,<br>Median of $\Phi_{1-3}$,<br>Mean of $\Phi_{1-3}$,<br>Mean of $\Phi_3$, Median<br>of $\Phi_3$, Mean of<br>Means $\Phi_{1-4}$, Mean<br>of GLM BLUE<br>Fitted $\Phi_{1-4}$ | 0.10 |
| Chr02 | 38310584 | Mean of $\Phi_{1-4}$ | Median of $\Phi_{1-3}$,<br>Median of $\Phi_3$,<br>Mean of $\Phi_2$, Mean<br>of GLM BLUE<br>Fitted $\Phi_{1-4}$ | 0.10 |
| Chr02 | 53403506 | Mean of $\Phi_{1-4}$ | Median of $\Phi_{1-3}$,<br>Mean of Means<br>$\Phi_{1-4}$, Mean of<br>GLM BLUE Fitted<br>$\Phi_{1-4}$ | 0.10 |

**Supplementary Table 3:** Identified genetic markers for Median Φ_1*−*4_ are within 100kb of multiple gene models.

| Gene model | Genetic marker | Distance from genetic marker (kb) | LD ($R^2$) | Functional annotation |
| --- | --- | --- | --- | --- |
| Sobic.005G086600 | Chr05:12,109,370 | 76.6 | 0.01 – 0.06 | O-methyltransferase |
| Sobic.005G086650 | Chr05:12,109,370 | 55.6 | 0.00 – 0.11 | None available |
| Sobic.005G086700 | Chr05:12,109,370 | 24.5 | 0.01 – 0.18 | Zinc finger transcription factor |
| Sobic.005G086801 | Chr05:12,109,370 | 51.1 | 0.00 – 0.11 | None available |
| Sobic.005G086900 | Chr05:12,109,370 | 64.9 | 0.07 – 0.09 | None available |
| Sobic.005G175200 | Chr05:65,733,791 | 46.5 | 0.01 – 0.01 | Serine/threonine protein kinase |
| Sobic.005G175300 | Chr05:65,733,791 | 33.6 | 0.00 – 0.38 | None available |
| Sobic.005G175400 | Chr05:65,733,791 | 16.4 | 0.00 – 0.69 | None available |
| Sobic.005G175500 | Chr05:65,733,791 | 0.0 | 0.00 – 1.00 | Protein tyrosine kinase related to salt stress/antifungal response |
| Sobic.005G175600 | Chr05:65,733,791 | 4.3 | 0.00 – 0.00 | Protein tyrosine kinase related to salt stress/antifungal response |
| Sobic.005G175700 | Chr05:65,733,791 | 18.1 | 0.01 – 0.45 | Ubiquitin-activating enzyme E1 |
| Sobic.005G175800 | Chr05:65,733,791 | 22.6 | 0.02 – 0.18 | Calcium binding protein CML30-related |
| Sobic.005G175900 | Chr05:65,733,791 | 26.9 | 0.01 – 0.45 | None available |
| Sobic.005G176000 | Chr05:65,733,791 | 37.8 | 0.00 – 0.43 | Membrane-associated zinc finger protein |
| Sobic.005G176100 | Chr05:65,733,791 | 47.2 | 0.01 – 0.11 | Mannose-6-phosphate isomerase |
| Sobic.005G176300 | Chr05:65,733,791 | 82.3 | 0.01 – 0.22 | Leucine-rich repeat-containing protein |
| Sobic.006G061000 | Chr06:41,390,777 | 32.4 | 0.68 – 0.76 | PPR repeat family |
| Sobic.006G061100 | Chr06:41,390,777 | 18.8 | 0.61 – 0.71 | AMP-activated protein kinase |
| Sobic.006G061201 | Chr06:41,390,777 | 57.8 | 0.12 – 0.12 | None available |
| Sobic.006G061300 | Chr06:41,390,777 | 65.7 | 0.18 – 0.20 | Osmotic stress potassium transporter |

**Supplementary Figure 1:**
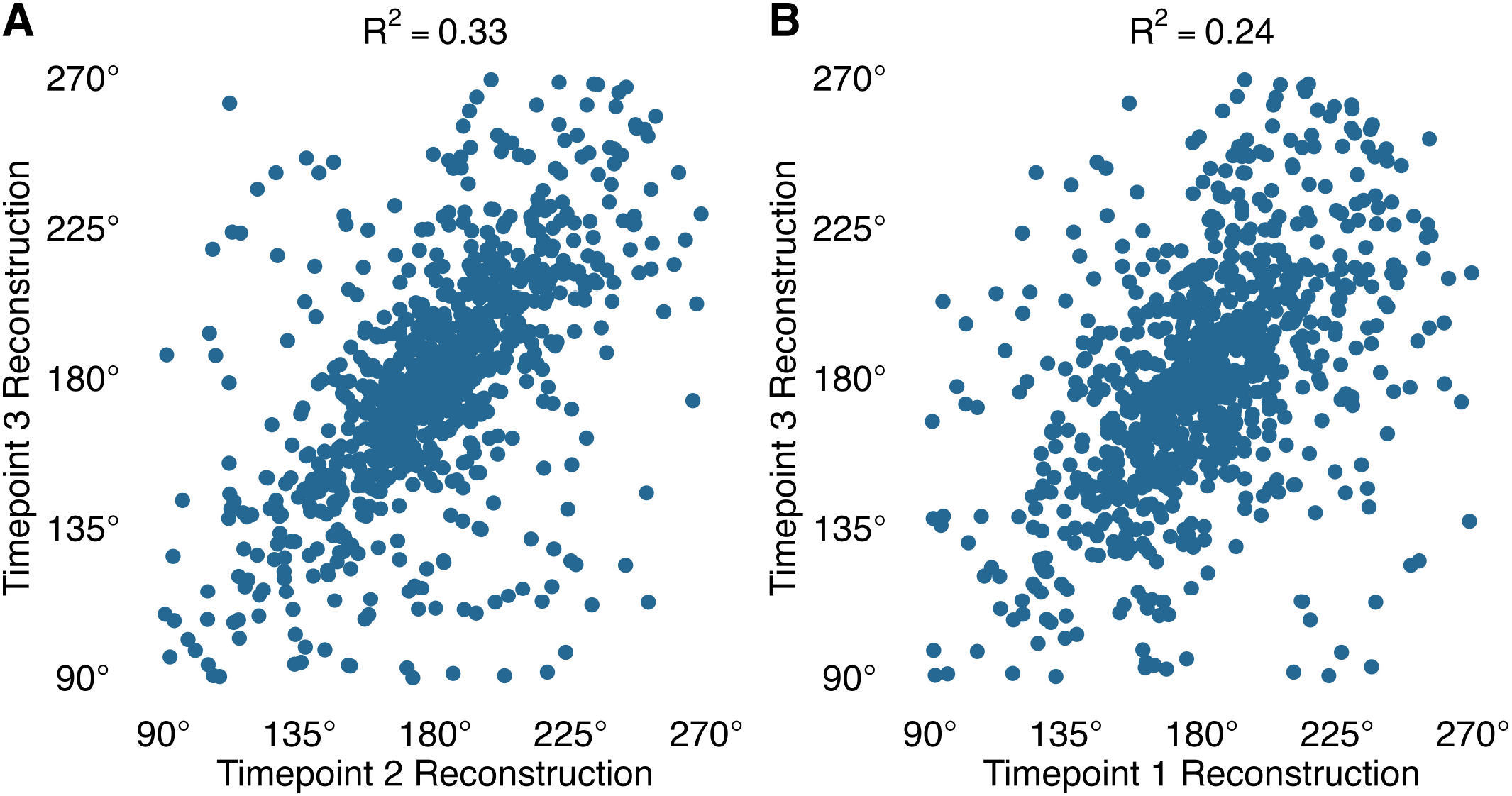
Correlation between phyllotaxic angles from reconstructions of the same plant decreases as time between imaging dates increases. **A)** Correlation of lower five *φ* values measured by 3D reconstructions of the sorghum plants when imaged on April 13, 2018 (Timepoint 2) and April 16, 2018 (Timepoint 3) after removing *φ* values less than 90° or greater than 270°, with an *R*^2^ value of 0.3304 (*n* = 820 angles). In cases when the measurements for a single plant were negatively correlated between the two days, the values from the Timepoint 3 reconstruction were transformed to their conjugate angles. **B)** Correlation of lower five *φ* values measured by 3D reconstructions of the sorghum plants when imaged on April 11, 2018 (Timepoint 1) and April 16, 2018 (Timepoint 3) after removing *φ* values less than 90° or greater than 270°, with an *R*^2^ value of 0.2351 (*n* = 820 angles). In cases when the measurements for a single plant were negatively correlated between the two days, the values from the Timepoint 3 reconstruction were transformed to their conjugate angles.

**Supplementary Figure 2:**
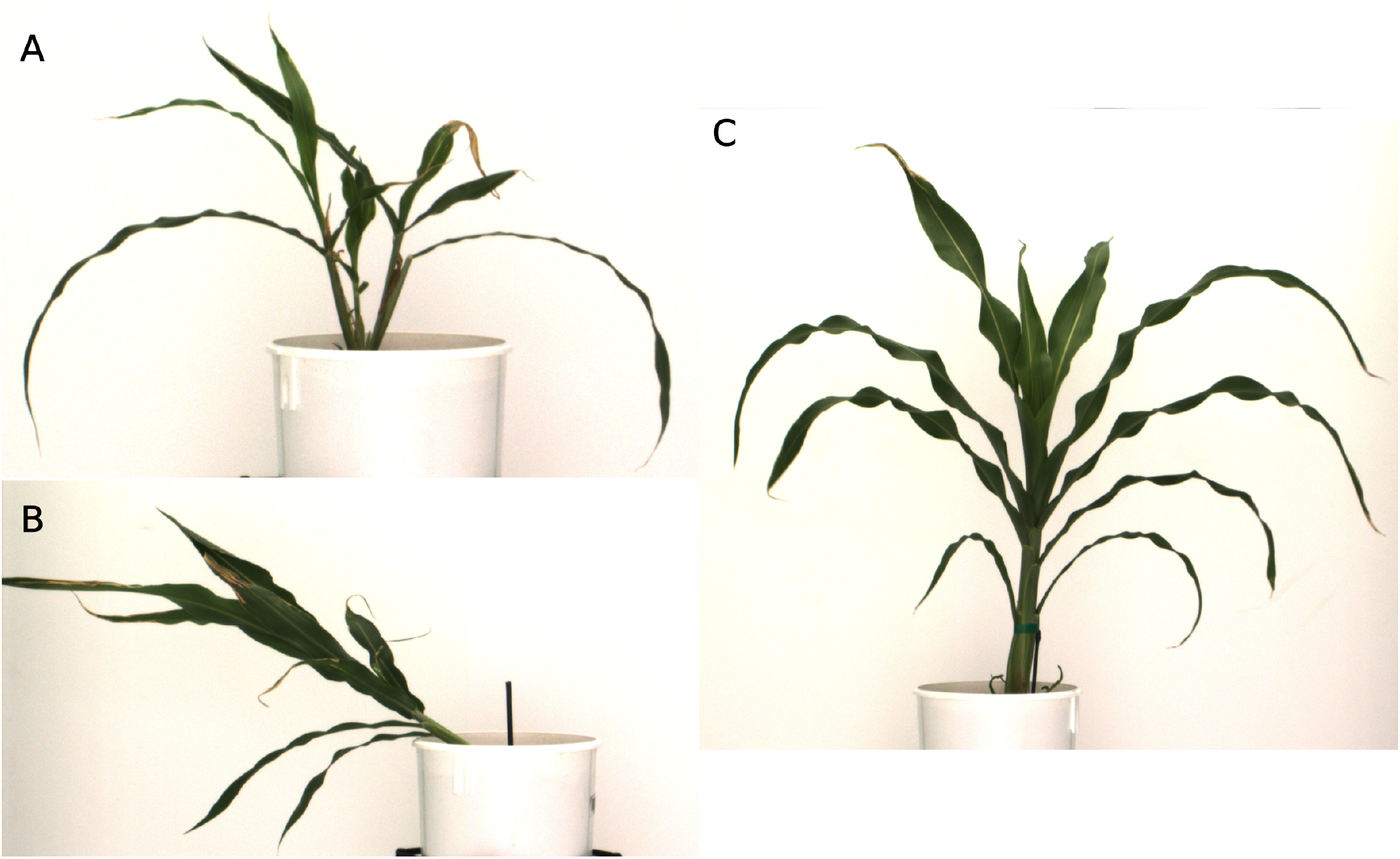
Tillers, lodging, or early leaf senescence in sorghum plants can cause extreme phyllotaxic values in reconstruction measurements. **A)** A sorghum plant with large tillers. This can cause extreme phyllotaxy values when measured via 3D reconstruction. A lodged sorghum plant. Lodged plants may also present as having extreme phyllotaxy values. A plant where the third true leaf has senesced before the first and second true leaves, resulting in an extreme phyllotaxic angle being measured as *φ*_2_ between the second true leaf and the fourth true leaf.

**Supplementary Figure 3:**
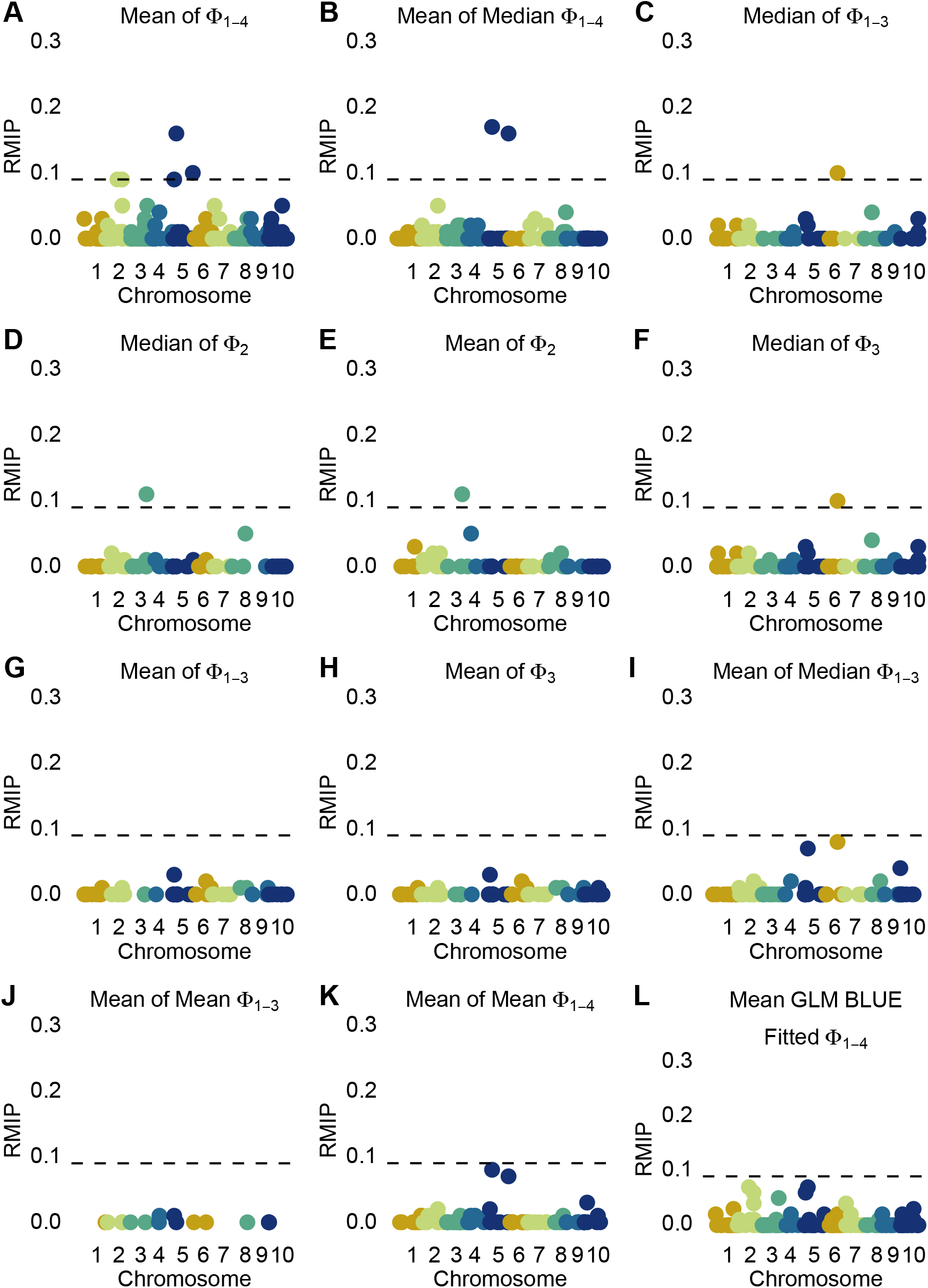
Resampling FarmCPU GWAS detects stable marker-trait associations for multiple quantitative summaries of lower-canopy phyllotaxy. **A – L)** Manhattan plot of marker-trait associations for each quantitative summary of lower canopy phyllotaxy with a broad-sense heritability *≥* 0.20.

