## Supplementary material for "3D Reconstruction Enables High-Throughput Phenotyping and Quantitative Genetic Analysis of Phyllotaxy": Figure 1 Publication License

49 Spadina Ave. Suite 200
Toronto ON M5V 2J1 Canada
www.biorender.com

### Confirmation of Publication and Licensing Rights - Open Access

**October 3rd, 2024**

| | |
|---|---|
| ***Subscription Type:*** | *Institution - Academic* |
| ***Agreement number:*** | *HM27DRKES8* |
| ***Publisher Name:*** | *bioRxiv* |
| ***Figure Title:*** | *Phyllotaxy is the arrangement of the leaves around the stem in the xy plane.* |
| ***Citation to Use:*** | *Created in BioRender. Torres, V. (2024) BioRender.com/a16d728* |

To whom this may concern,

This document ("Confirmation") hereby confirms that Science Suite Inc. dba BioRender ("BioRender") has granted the following BioRender user: Vladimir Torres ("User") a BioRender Academic Publication License in accordance with BioRender's Terms of Service and Academic License Terms ("License Terms") to permit such User to do the following on the condition that all requirements in this Confirmation are met:

1) publish their Completed Graphics created in the BioRender Services containing both User Content and BioRender Content (as both are defined in the License Terms) in publications (journals, textbooks, websites, etc.); and
2) sublicense such Completed Graphics under "open access" publication sublicensing models such as CC-BY 4.0 and more restrictive models, so long as the conditions set forth herein are fully met.

Requirements of User:

1) All Completed Graphics to be published in any publication (journals, textbooks, websites, etc.) must be accompanied by the following citation either as a caption, footnote or reference for each figure that includes a Completed Graphic:
"Created in BioRender. Torres, V. (2024) BioRender.com/a16d728 ".
2) All terms of the License Terms including all Prohibited Uses are fully complied with. E.g. For Academic License Users, no commercial uses (beyond publication in journals, textbooks or websites) are permitted without obtaining or switching to a BioRender Industry Plan.
3) A Reader (defined below) may request that the User allow their figure to be a public template for Readers to view, copy, and modify the figure. It is up to the User to determine what level of access to grant.

Open-Access Journal Readers:

Open-Access journal readers ("Reader") who wish to view and/or re-use a particular Completed Graphic in an Open-Access journal subject to CC-BY sublicensing may do so by clicking on the URL link in the applicable citation for the subject Completed Graphic.

The re-use/modification options below are available after the Reader requests the User to adapt their figure as a BioRender template and the User has granted such access.

1) View-Only/Free Plan Use: A Reader who wishes to only view the Completed Graphic may do so in the BioRender Services as either a BioRender Free Plan user or simply as a viewer. By becoming a BioRender Free Plan user, the Reader may view, modify and re-use the Completed Graphic as permitted under BioRender's Basic License Terms (e.g. personal use only, no publishing or commercial use permitted).
2) Re-Use/Publish with No Modifications: For any re-use and re-publication of a Completed Graphic with no modification(s) to the Completed Graphic made by the Reader, a Reader may do so by citing the original author using the citation noted above with the Completed Graphic. The Reader must also comply with the underlying License Terms which apply to the Completed Graphic as noted above (e.g. no commercial use for Academic License).
3) Re-Use/Publish with Modifications: For any re-use and re-publication of a Completed Graphic with a modification(s) made by the Reader, the Reader may do so by becoming a BioRender user themselves under either an Academic or Industry Plan, citing the original author using the citation noted above with the Completed Graphic and complying with the applicable License Terms.

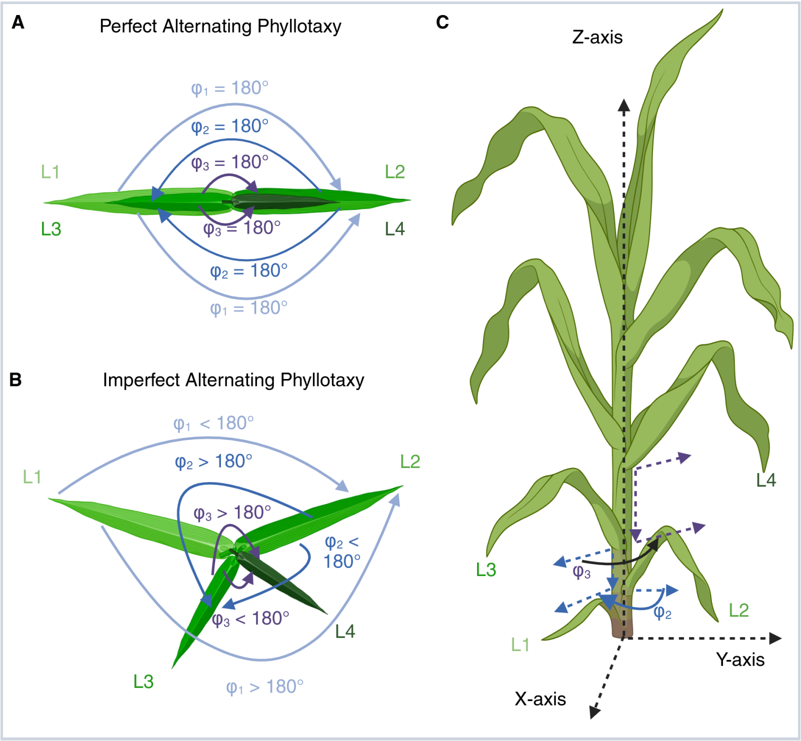


*For any questions regarding this document, or other questions about publishing with BioRender, please refer to our BioRender Publication Guide, or contact BioRender Support at.*
